# Unexpected functional role of the transactivation domain for nuclear import of STAT5

**DOI:** 10.64898/2025.12.11.693681

**Authors:** Sabrina Ernst, Merith Borgen-Möller, Andrea Küster, Henning Schurse, Gerhard Müller-Newen

## Abstract

Signal transducer and activator of transcription 5 (STAT5) is a key transcriptional regulator acting downstream of hematopoietic cytokines and hormones, such as erythropoietin (Epo), thrombopoietin or prolactin. STAT5-mediated gene regulation involves tyrosine phosphorylation at cytokine receptors and subsequent nuclear import. We studied STAT5 nucleocytoplasmic shuttling via live-cell imaging of fluorescent mutants in STAT5^-/-^ HeLa EpoR cells. Unexpectedly, STAT5 mutants lacking the transactivation domain (TAD) were retained in the cytoplasm following Epo stimulation. Building upon this, we identified a 12-amino-acid stretch in the TAD sufficient to restore nuclear translocation. Further analysis revealed two residues within this 12-amino-acid stretch, D754 and D758, to be essential for nuclear import of phosphorylated full-length STAT5. Importantly, a single intact TAD in the STAT5 dimer is sufficient for nuclear import. Our findings reveal a unique role of the TAD in STAT5 nuclear trafficking distinct from other STATs, providing new mechanistic insight and potential targets for therapeutic intervention in STAT5-driven disease such as myeloproliferative neoplasms and leukemia.

## Introduction

Signal transducer and activator of transcription (STAT) proteins are latent transcription factors that function as molecular messengers, transmitting extracellular cues to the nucleus, where they modulate gene expression^1^. STATs are activated at cytokine or growth factor receptors through phosphorylation of a critical tyrosine residue. Phosphorylated STATs (pSTATs) dimerize via reciprocal phosphotyrosine-SH2 domain interactions, resulting in nuclear accumulation of the dimers followed by DNA-binding and finally induction of gene transcription^2, 3^. Dysregulated STAT activation contributes to numerous pathophysiological processes including chronic inflammation and cancer^4^.

While nuclear import of pSTAT dimers is essential for STAT function, the underlying molecular mechanisms vary among family members and are incompletely defined^5^. Best characterized is STAT1, which mediates interferon-induced antiviral responses^6^. The pSTAT1 dimer binds importin-α5 in an unconventional manner^7, 8^ that does not dependent on a classical nuclear localization signal (NLS) but instead utilizes a dimer-specific NLS (dsNLS) comprising residues within the DNA-binding (DBD)^9^ and SH2 domains^10^. Other STATs pass the nuclear pore complex (NPC) also after binding to carriers of the importin-α family. However, the molecular mechanisms underlying these processes are less well defined. A classical, transferable NLS has so far not been unequivocally identified in any of the STAT proteins. Residues involved in the dsNLS have been mapped primarily to the N-terminal domain (NTD), coiled-coil domain (CCD) and DBD^5^.

STAT5, activated downstream of erythropoietin (Epo) and other hematopoietic cytokines^11^, exerts essential functions in hematopoiesis. Persistent or dysregulated STAT5 activation drives hematologic malignancies, including myeloproliferative neoplasms (MPN) and leukemia^12^. In the endocrine system, STAT5 mediates growth hormone and prolactin signaling to regulate body growth and mammary gland differentiation^13, 14^. STAT5 exists as two closely related isoforms, STAT5A and STAT5B, which share approximately 95% amino acid identity but exhibit nonredundant functions due to tissue-specific expression patterns and differential recruitment of cofactors mediated primarily by their most divergent regions, the transactivation domains (TADs)^15^.

Like STAT1 and STAT3^9, 16^, latent STAT5 continuously shuttles between the cytoplasm and the nucleus maintaining a dynamic steady-state distribution^17^. Nuclear export of STATs involves exportin-1 (Crm1), as evidenced by leptomycin B-induced partial nuclear accumulation^17–19^. Consistent with STAT1 and STAT3^20, 21^, the NTD is essential for nuclear import of pSTAT5 as has been shown for STAT5B^17^. In addition, regions in the CCD have been described to be required for nuclear import of pSTAT5^19, 22^. However, evidence for association with α-importins remains inconsistent. Interaction of STAT5 with importin-α3 has been reported^19^ while others failed to detect interaction with any of the relevant α-importins^23^.

Unexpectedly, we observed that a STAT5 deletion mutant lacking the TAD displayed cytoplasmic rather than nuclear accumulation after Epo stimulation. Building upon this observation, we identified two amino acid residues within the TAD as critical determinants of pSTAT5 nuclear import. Among STAT family members, involvement of the TAD in nuclear accumulation appears unique to STAT5 and may represent a selective target for therapeutic intervention.

## Results

### 1. The transactivation domain is essential for nuclear accumulation of STAT5 in response to activation by Epo or the BCR-ABL oncogene

Upon appropriate cytokine stimulation of cells, STAT transcription factors accumulate in the nucleus^5, 24, 25^. Accordingly, a fusion protein of STAT5A and monomeric enhanced YFP (STAT5A-meYFP) translocated to the nucleus upon erythropoietin (Epo) stimulation in HeLa cells stably expressing the Epo receptor (HeLa EpoR). Live-cell imaging shows a shift from an initially nearly equal cytoplasmic and nuclear distribution to a predominantly nuclear localization (Fig. 1A, upper panel).

**Figure 1:**
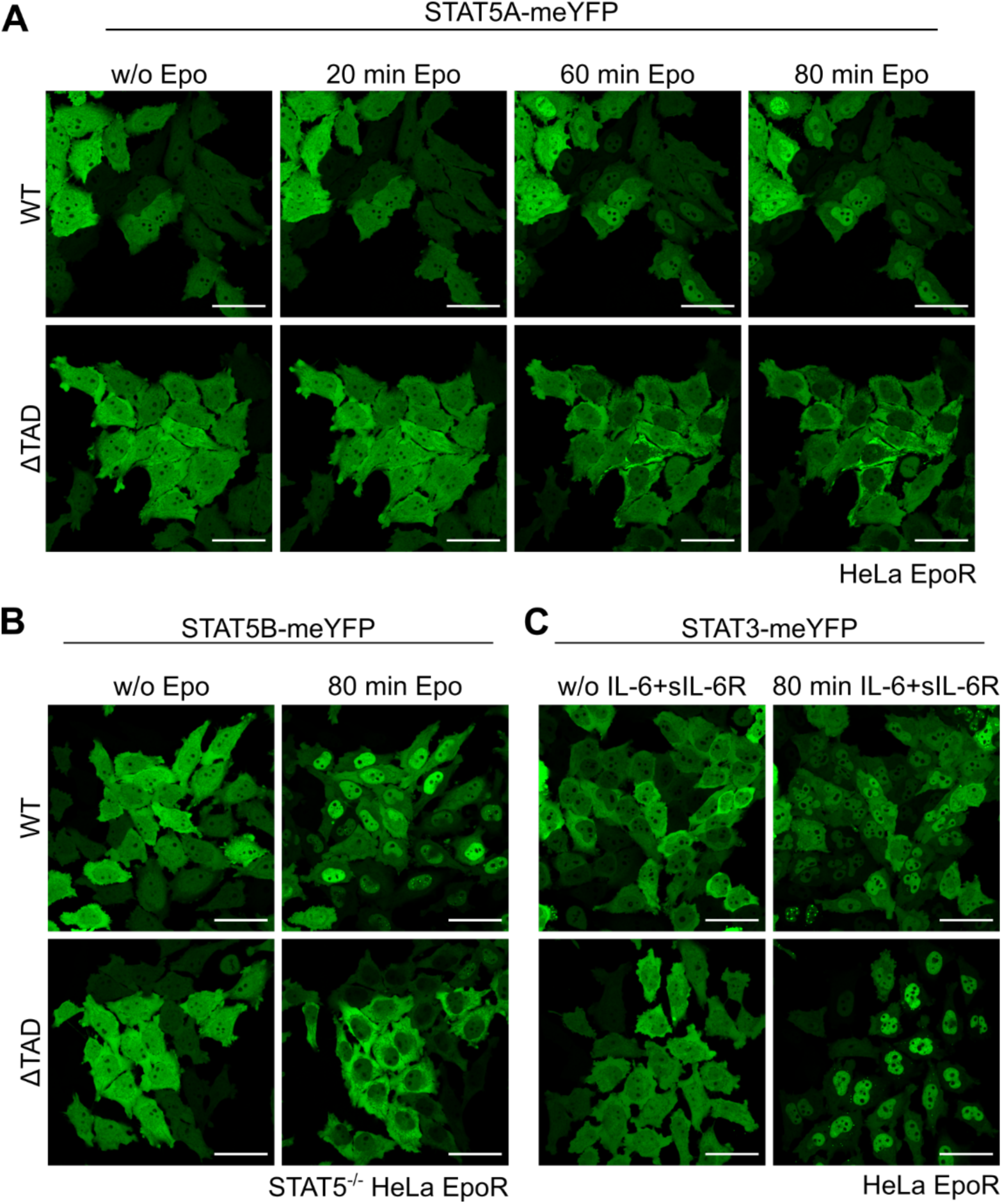
Cytokine-induced nucleocytoplasmic shuttling of STAT5A, STAT5B and STAT3 WT and ΔTAD deletion mutants. **(A)** HeLa T-REx cells were stably transduced with EpoR (HeLa EpoR) and subsequently with STAT5A-WT-meYFP or STAT5A-ΔTAD-meYFP. **(B)** HeLa EpoR cells deficient for STAT5A and STAT5B (STAT5^-/-^ HeLa EpoR) were transduced with STAT5B-WT-meYFP or STAT5B-ΔTAD-meYFP **(C)** HeLa EpoR cells were transduced with STAT3-WT-meYFP or STAT3-ΔTAD-meYFP. Cells were stimulated with **(A,B)** 1 U/ml Epo or **(C)** 20 ng/ml IL-6 and 1 µg/ml sIL-6R for the indicated times. meYFP fluorescence was detected via confocal live cell imaging. w/o, without; scale bars: 50 µm.

While characterizing several STAT5A deletion mutants, we found that a STAT5A-meYFP variant lacking the transactivation domain (STAT5A-ΔTAD-meYFP, truncated C-terminal to A717) failed to accumulate in the nucleus upon Epo stimulation. Instead, STAT5A-ΔTAD-meYFP showed cytoplasmic accumulation (Fig. 1A, lower panel). To investigate this unusual behavior in greater detail without interference from endogenous STAT5, HeLa EpoR cells lacking STAT5A and STAT5B were generated via CRISPR/Cas9 gene targeting (STAT5^-/-^ HeLa EpoR, Suppl. Fig. 1). In these cells, the corresponding STAT5B-ΔTAD-meYFP construct displayed the same cytoplasmic accumulation (Fig. 1B). In striking contrast, a STAT3-ΔTAD-meYFP construct exhibited even stronger nuclear accumulation upon IL-6 stimulation than the wild type (WT) STAT3-meYFP construct in HeLa EpoR cells (Fig. 1C).

To verify the paradoxical behavior of STAT5-ΔTAD, we conducted a series of control experiments. Cytoplasmic accumulation of STAT5A-ΔTAD-meYFP upon Epo stimulation was also observed in HEK cells transiently transfected with EpoR and the STAT5A constructs (Fig. 2A). Likewise, activation of STAT5A-ΔTAD-meYFP by the BCR-ABL oncogene let to cytoplasmic accumulation (Fig. 2B). Positioning the meYFP-tag at the N-terminus of STAT5A impaired nuclear accumulation of the WT protein but did not affect cytoplasmic accumulation of STAT5A-ΔTAD (Fig. 2C). Finally, after Epo stimulation, phosphorylated non-tagged STAT5A-ΔTAD was predominantly detected in the cytoplasm by immunofluorescence, whereas pSTAT5A WT was enriched in the nucleus (Fig. 2D). These results indicate that deletion of the TAD prevents nuclear import, leading to cytoplasmic accumulation of pSTAT5.

**Figure 2:**
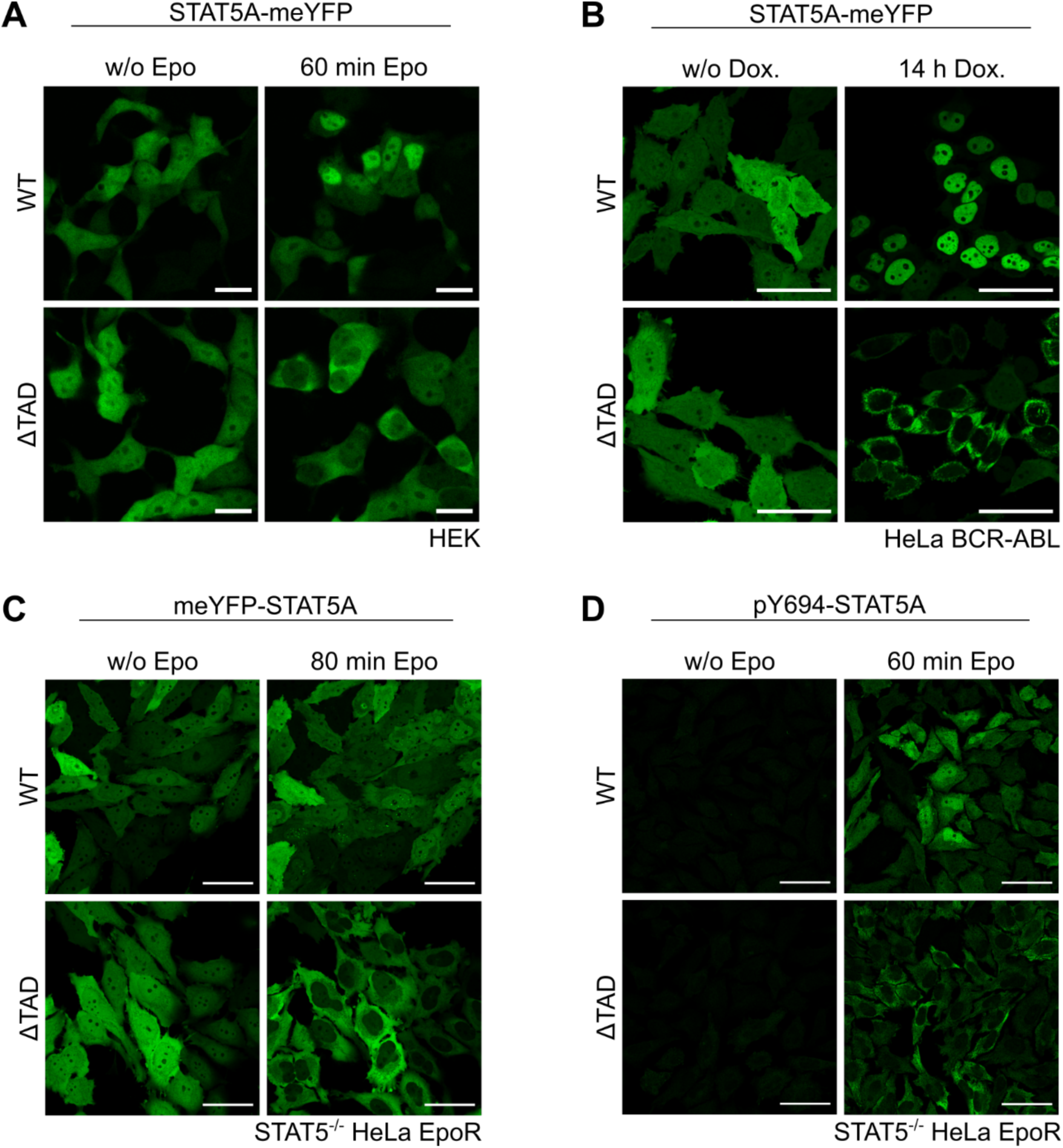
Nucleocytoplasmic shuttling of STAT5A WT and ΔTAD deletion mutant under different conditions. **(A)** HEK293T (HEK) cells were transiently cotransfected with EpoR and STAT5A-WT-meYFP or STAT5A-ΔTAD-meYFP and stimulated with Epo (1U/ml) as indicated. Note, that only those cells respond to Epo stimulation that were cotransfected with both Epo receptor and STAT5A constructs. Scale bars: 10 µm. **(B)** HeLa T-REx cells transduced with BCR-ABL (HeLa BCR-ABL) and subsequently with STAT5A-WT-meYFP or STAT5A-ΔTAD-meYFP were treated with 5 ng/ml doxycycline (Dox) as indicated. Scale bars: 50 µm. **(C)** STAT5^-/-^ HeLa EpoR cells transduced with meYFP-STAT5A-WT or meYFP-STAT5A-ΔTAD were stimulated with 1 U/ml Epo as indicated. meYFP fluorescence was detected via confocal live cell imaging. Scale bars: 50 µm. **(D)** STAT5^-/-^ HeLa EpoR cells transduced with STAT5A-WT or STAT5A-ΔTAD were stimulated with 1 U/ml Epo as indicated. Fixed cells were stained for pY694-STAT5A and analyzed by confocal microscopy. Scale bars: 50 µm.

### 2. Phosphorylation and dimerization are required for cytoplasmic accumulation of STAT5A-ΔTAD

Upon Epo pulse stimulation, STAT5A-ΔTAD-meYFP underwent phosphorylation and subsequent dephosphorylation with kinetics similar to those of STAT5A-meYFP (Fig. 3A). Deletion of the TAD did not affect STAT5A dimerization, as demonstrated by native PAGE (Fig. 3B). To assess whether phosphorylation of the critical tyrosine residue Y694 is required for cytoplasmic accumulation of STAT5A-ΔTAD, the STAT5A-ΔTAD-Y694F-meYFP mutant was generated (Fig. 3C, upper panel). This mutant did not change its subcellular localization upon Epo stimulation (Fig. 3C, lower panel). The dimerization-deficient STAT5A-ΔTAD-F706A-meYFP mutant^26^ was used to determine the requirement for dimerization in cytoplasmic accumulation of pSTAT5A-ΔTAD. Inhibition of phosphatases with sodium vanadate enhanced Epo-induced phosphorylation and dimerization (Fig. 3D) as well as cytoplasmic accumulation (Fig. 3E, lower panel) of STAT5A-ΔTAD-meYFP. In contrast, the dimerization-deficient STAT5A-ΔTAD-F706A-meYFP mutant displayed only minimal dimerization in the presence of sodium vanadate (Fig. 3D) and no substantial cytoplasmic accumulation upon Epo stimulation (Fig. 3E, upper and middle panels). These observations indicate that both tyrosine phosphorylation and dimerization are prerequisites for cytoplasmic accumulation of pSTAT5A-ΔTAD-meYFP.

**Figure 3:**
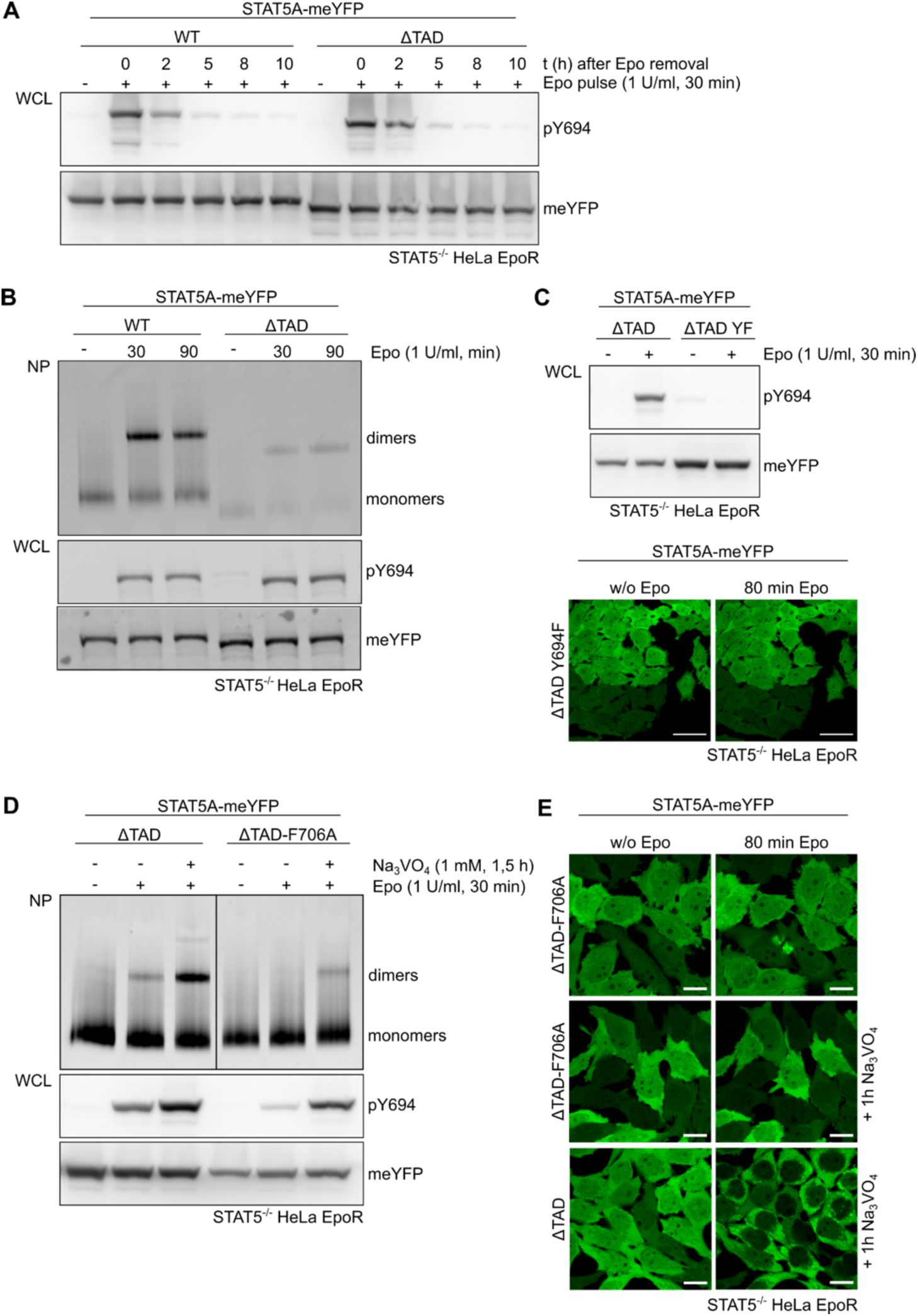
Phosphorylation and dimer formation of STAT5A mutants. STAT5^-/-^ HeLa EpoR cells transduced with STAT5A-WT-meYFP or STAT5A-ΔTAD-meYFP were **(A)** pulse-stimulated with 1 U/ml Epo for 30 minutes, whole cell lysates (WCL) were prepared after Epo removal at indicated time points and analyzed by immunoblotting with the indicated antibodies or were **(B)** stimulated with 1 U/ml Epo for the indicated time points and WCL analyzed by native PAGE (NP) or immunoblotting with the indicated antibodies. **(C)** Upper panel: STAT5^-/-^ HeLa EpoR cells transduced with STAT5A-ΔTAD-meYFP or STAT5A-ΔTAD-Y694F-meYFP were stimulated with 1 U/ml Epo as indicated. WCL were analyzed by immunoblotting with the indicated antibodies. Lower panel: Additionally, meYFP fluorescence was detected via confocal live cell imaging as indicated. Scale bars: 50 µm. **(D)** STAT5^-/-^ HeLa EpoR cells transduced with STAT5A-ΔTAD-meYFP or STAT5A-ΔTAD-F706A-meYFP were treated with 1 mM Na_3_VO_4_ and/or 1 U/ml Epo as indicated and WCL were analyzed by native PAGE (NP) or immunoblotting with the indicated antibodies. **(E)** Cell lines transduced as described in (D) were treated with 1 mM Na_3_VO_4_ and/or 1 U/ml Epo as indicated. meYFP fluorescence was detected via confocal live cell imaging. w/o, without; scale bars: 20 µm.

### 3. Apparent cytoplasmic accumulation of STAT5A-ΔTAD does not involve Crm1-mediated export, proteasomal degradation or lysine acetylation

Treatment of cells with the Crm1 (exportin-1) inhibitor leptomycin B has been reported to cause partial nuclear accumulation of STAT5, supporting the notion of persistent nucleocytoplasmic shuttling of latent STAT5^17^. Consistent with these findings, leptomycin B treatment of STAT5^-/-^ HeLa EpoR cells induced a decent shift of STAT5A-meYFP into the nuclear compartment (Suppl. Fig. 2A, left panel).

However, leptomycin B did not affect the Epo-induced cytoplasmic accumulation of STAT5A-ΔTAD-meYFP (Suppl. Fig. 2A, right panel) indicating that cytoplasmic accumulation does not result from increased Crm1-mediated nuclear export.

Proteasomal degradation of nuclear STAT5 has been reported previously^27^. Pretreatment of HeLa STAT5^-/-^ EpoR cells with the proteasome inhibitor MG132 did not affect the subcellular distribution of STAT5A-ΔTAD-meYFP after Epo stimulation (Suppl. Fig. 2B), demonstrating that the apparent cytoplasmic accumulation is not due to Epo-induced nuclear degradation of STAT5A-ΔTAD.

The best characterized function of the STAT5 TAD is its interaction with the histone acetyltransferases p300/CBP^28^ which also catalyze lysine acetylation of STAT5. A STAT5A mutant carrying arginine substitutions at the reported lysine acetylation sites K696 and K700 (KK/RR)^29^ dimerized and accumulated in the nucleus in response to Epo similarly to WT (Suppl. Fig. 3). Thus, the cytoplasmic accumulation of STAT5A-ΔTAD is not caused by loss of p300/CBP-mediated lysine acetylation.

### 4. Specific requirement of the STAT5A TAD for nuclear translocation

Deletion of the TAD in STAT5A and STAT5B resulted in cytoplasmic accumulation, whereas deletion of the TAD in STAT3 caused enhanced nuclear accumulation after stimulation (Fig. 1), suggesting distinct roles of the STAT5 and STAT3 TADs in nucleocytoplasmic shuttling. To investigate this difference, chimeric constructs were generated in which the TAD of STAT5 was replaced by that of STAT3 and vice versa (Fig. 4A). Both chimeras, STAT5A-3TAD-meYFP and STAT3-5ATAD-meYFP, underwent tyrosine phosphorylation and dimerization upon stimulation with their respective cytokines (Fig. 4B and C). Notably, STAT3-5ATAD-meYFP showed enhanced phosphorylation and dimerization compared to STAT3-meYFP. Upon Epo stimulation, STAT5A-3TAD-meYFP failed to accumulate in the nucleus and instead localized predominantly to the cytoplasm, similar to STAT5A-ΔTAD-meYFP. (Fig. 4D, upper panel). In contrast, STAT3-5ATAD-meYFP not only accumulated in the nucleus but also formed punctate structures (Fig. 4D, lower panel) which may indicate liquid-liquid phase separation (LLPS). These data demonstrate that the TAD of STAT3 cannot compensate for the nuclear translocation function of the STAT5A TAD. Thus, the TAD of STAT5 plays a specific and non-redundant role in mediating nuclear import of phosphorylated STAT5 dimers.

**Figure 4:**
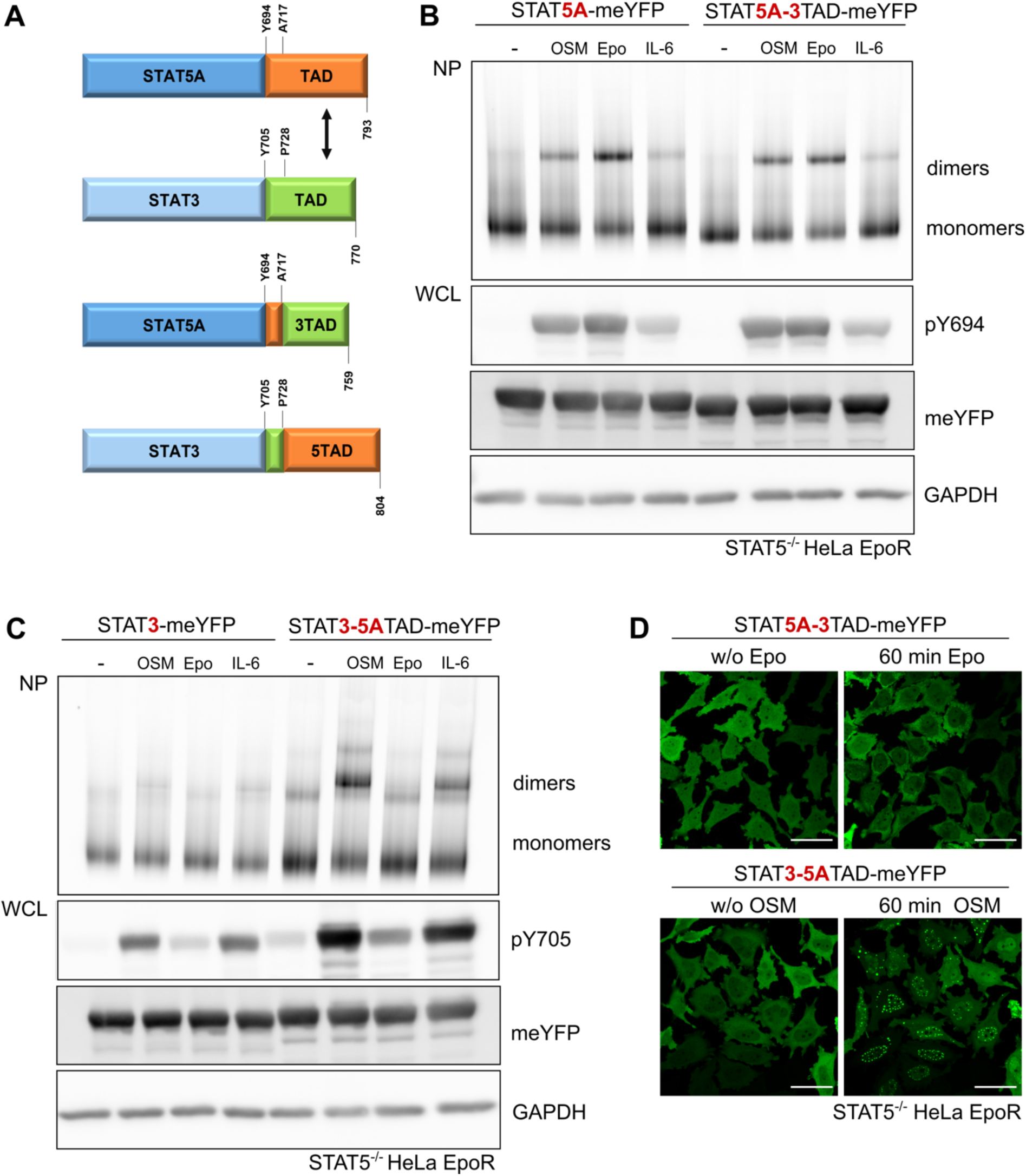
Dimerization and subcellular localization of STAT5A/STAT3 TAD chimeras. **(A)** Scheme of STAT5A and STAT3 TAD chimeras (dark blue: STAT5A; orange: TAD of STAT5A; light blue: STAT3; green: TAD of STAT3). **(B)** STAT5^-/-^ HeLa EpoR cells transduced with STAT5A-meYFP or STAT5A-3TAD-meYFP or **(C)** STAT3-WT-meYFP or STAT3-5ATAD-meYFP were treated with either 10 ng/ml OSM, 2 U/ml Epo or 20 ng/ml IL-6 and 0.5 µg/ml sIL-6R as indicated. Whole cell lysates (WCL) were analyzed by native PAGE (NP) or immunoblotting with the indicated antibodies. **(D)** Upper panel: STAT5^-/-^ HeLa EpoR cells transduced with STAT5A-3TAD-meYFP were stimulated with 2 U/ml Epo as indicated. Lower panel: STAT5^-/-^ HeLa EpoR cells transduced with STAT3-5ATAD-meYFP were stimulated with 10 ng/ml OSM as indicated. meYFP fluorescence was detected via confocal live cell imaging. w/o, without; scale bars: 50 µm.

### 5. A twelve-amino-acid stretch (751-762) is sufficient to restore nuclear accumulation of STAT5-ΔTAD

To further explore the function of the TAD in STAT5 nucleocytoplasmic shuttling, additional deletion mutants were generated to narrow down the critical amino acid region governing Epo-mediated nuclear versus cytoplasmic accumulation. Previous studies identified the region between F751 and H762 as important for STAT5A turnover and transcriptional activity^30, 31^. Accordingly, in addition to STAT5A-ΔTAD-meYFP (truncated at A717), the deletion mutants STAT5A-Δ750-meYFP (truncated behind E750), STAT5A-Δ756-meYFP (truncated behind S756), and STAT5A-Δ762-meYFP (truncated behind H762) were analyzed. Upon Epo stimulation of transduced STAT5^-/-^ HeLa EpoR cells, all mutants underwent tyrosine phosphorylation and formed dimers (Fig. 5A). However, whereas STAT5A-Δ762-meYFP translocated efficiently into the nucleus, STAT5A-Δ750-meYFP accumulated in the cytoplasm. STAT5A-Δ756-meYFP showed an intermediate phenotype, with no detectable nuclear accumulation and less pronounced nuclear clearance compared to STAT5A-Δ750-meYFP (Fig. 5B). These results demonstrate that the twelve amino acids spanning F751 to H762, previously implicated in transactivation and proteasomal turnover, critically affect the subcellular localization of activated STAT5A.

**Figure 5:**
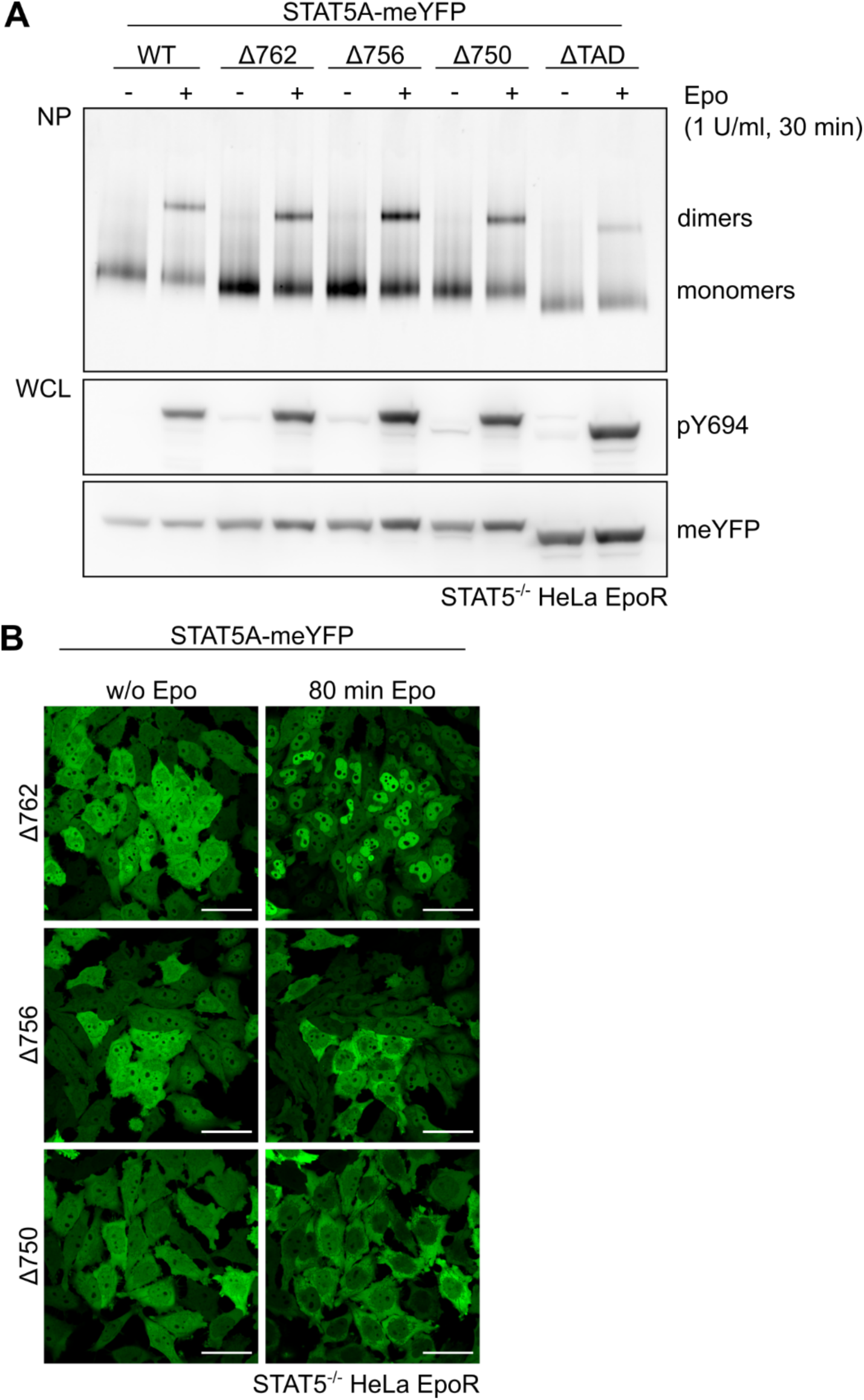
Dimerization and subcellular localization of STAT5A TAD deletion mutants. **(A)** STAT5^-/-^ HeLa EpoR cells transduced with STAT5A-WT-meYFP, STAT5A-Δ762-meYFP, STAT5A-Δ756-meYFP, STAT5A-Δ750-meYFP or STAT5A-ΔTAD-meYFP were stimulated with 1 U/ml Epo as indicated. Whole cell lysates (WCL) were analyzed by native PAGE (NP) or immunoblotting with the indicated antibodies. **(B)** STAT5^-/-^ HeLa EpoR cells transduced with STAT5A-Δ762-meYFP, STAT5A-Δ756-meYFP or STAT5A-Δ750-meYFP were stimulated with 1 U/ml Epo as indicated. meYFP fluorescence was detected via confocal live cell imaging. w/o, without; scale bars: 50 µm.

To determine whether this twelve-amino-acid stretch can restore nuclear accumulation of STAT5A-ΔTAD-meYFP and the STAT5A-3TAD-meYFP chimera, rescue mutants incorporating the F751-H762 sequence were generated (Fig. 6A). After Epo and oncostatin M (OSM) stimulation of transduced STAT5^-/-^ HeLa EpoR cells, both rescue mutants, STAT5A-ΔTAD+12aa-meYFP and STAT5A-3TAD+12aa-meYFP, were tyrosine phosphorylated and dimerized (Fig. 6B). Strikingly, these mutants translocated into the nucleus upon stimulation, like STAT5A-WT-meYFP (Fig. 6C).

**Figure 6:**
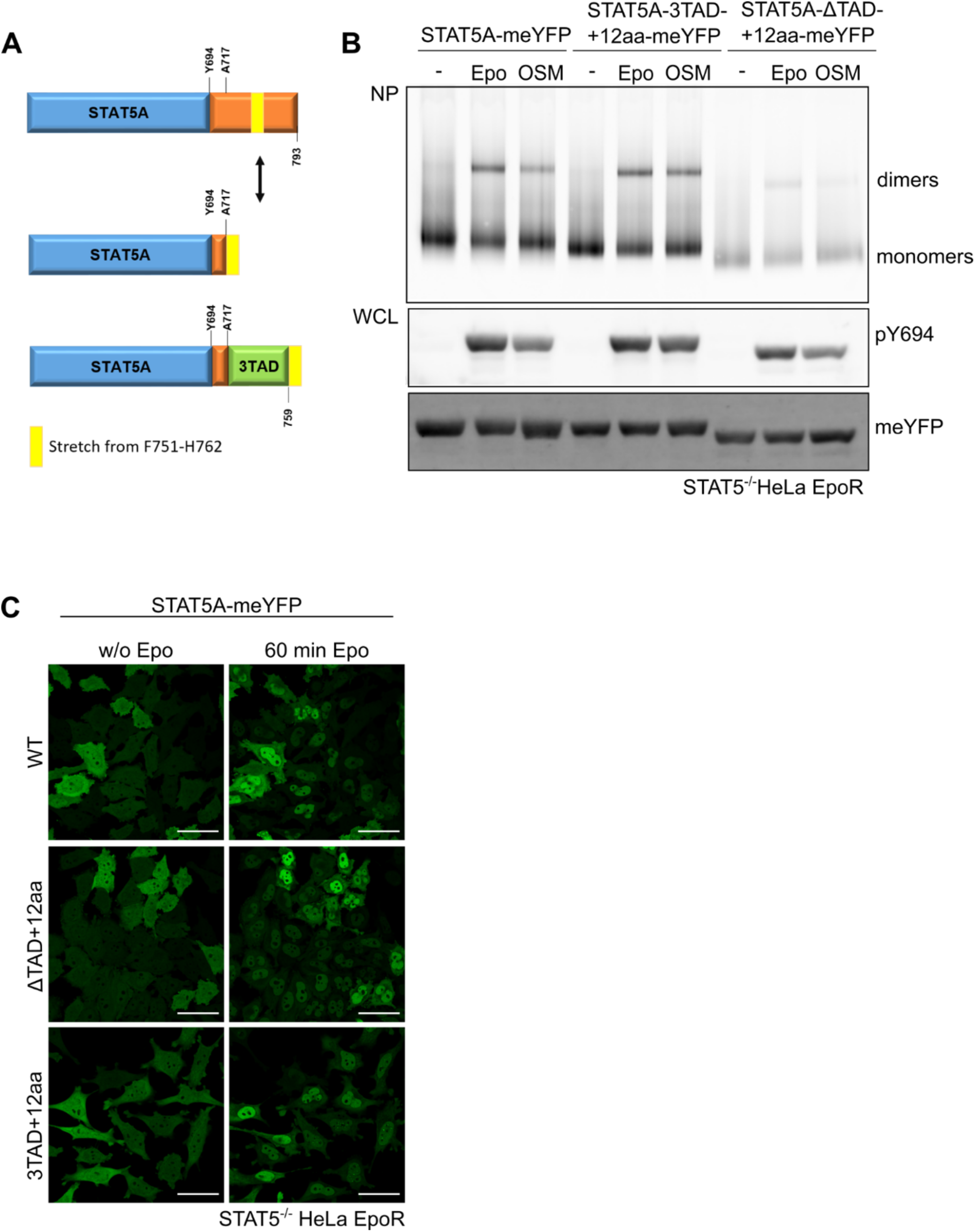
Dimerization and subcellular localization of STAT5A rescue mutants. **(A)** Scheme of STAT5A-ΔTAD and STAT5A-3TAD rescue mutants. Dark blue: STAT5A; orange: TAD of STAT5A; green: TAD of STAT3, yellow: stretch of twelve amino acids from F751 to H762. **(B)** STAT5^-/-^ HeLa EpoR cells transduced with STAT5A-WT-meYFP, STAT5A-ΔTAD+12aa-meYFP or STAT5A-3TAD+12aa-meYFP were stimulated with either 2 U/ml Epo or 10 ng/ml OSM as indicated. Whole cell lysates (WCL) were analyzed by native PAGE (NP) or immunoblotting with the indicated antibodies. **(C)** Cell lines described in (B) were stimulated with 2 U/ml Epo as indicated. meYFP fluorescence was detected via confocal live cell imaging. w/o, without; scale bars: 50 µm.

### 6. Aspartic acids 754 and 758 are essential for nuclear accumulation of STAT5A

The functional significance of subregions within the twelve-amino-acid stretch was investigated by alanine triplet scanning mutagenesis within full-length STAT5A, which identified the central six residues D754-E-S-M-D-V759 as critical (Suppl. Fig. 4). In this sequence, a clustering of acidic amino acids stands out, along with a potential phosphorylation site at S756, which is replaced by threonine in STAT5B (Suppl. Fig. 5A). Mutation of S756 to alanine had no effect on dimerization and nuclear import of pSTAT5A (Suppl. Fig. 5B and C). However, mutation of the acidic residues D754 and D758 to alanine (STAT5A-2Dto2A) resulted in impaired nuclear translocation and cytoplasmic retention of STAT5A (Fig. 7A, upper panel) without affecting tyrosine phosphorylation and dimerization (Fig. 7B). These results highlight D754 and D758 as key residues for nuclear import of STAT5A. However, add back of these two aspartic residues in the twelve-amino-acid stretch, where the other amino acids were mutated to alanine (STAT5A-10Aand2D), was insufficient to restore nuclear translocation (Fig. 7A, lower panel). This finding indicates that although these acidic residues are necessary, other residues within the stretch also contribute to efficient nuclear accumulation of STAT5A.

**Figure 7:**
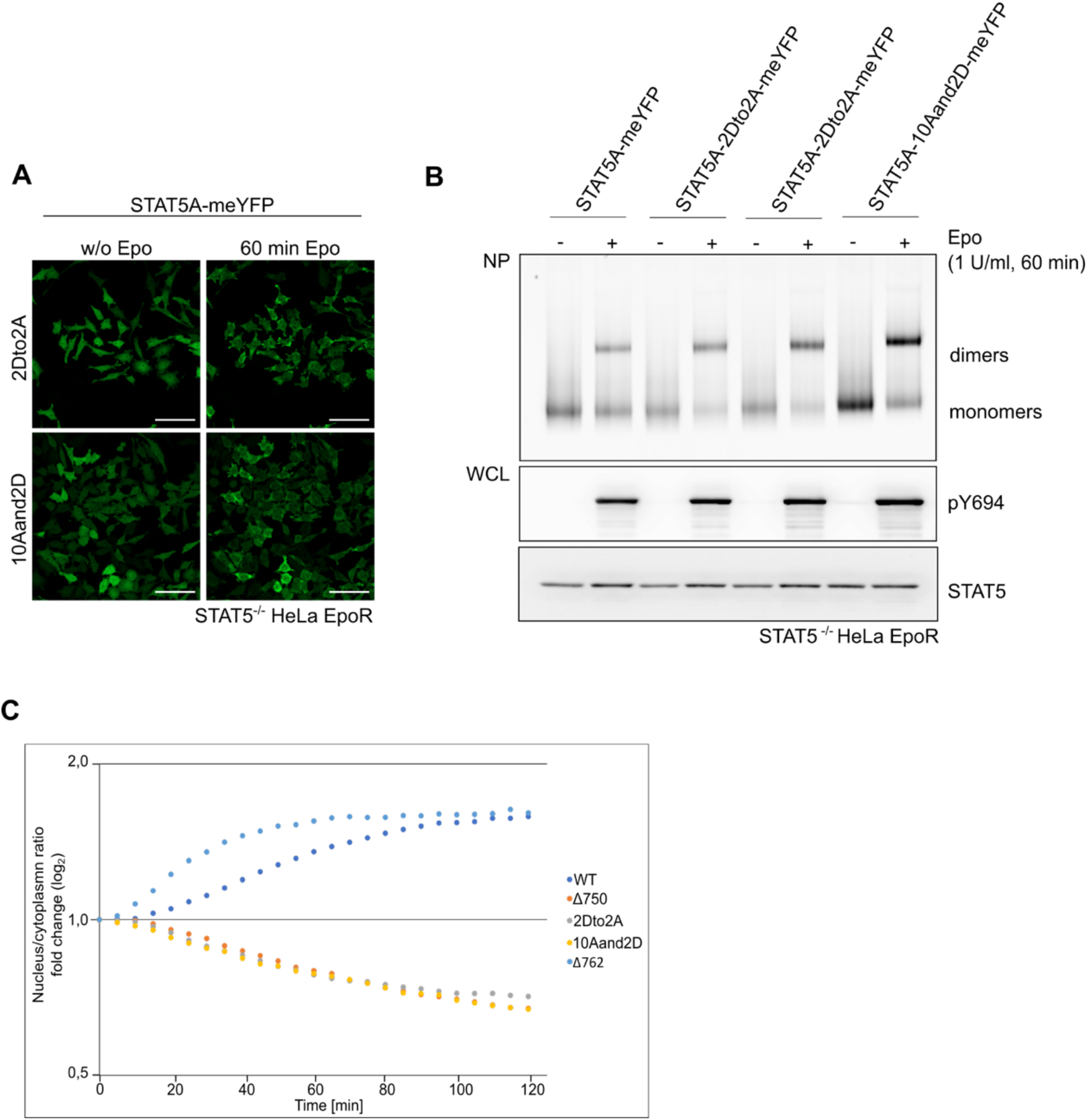
Dimerization and subcellular localization of STAT5A-2Dto2A and STAT5A-10Aand2D mutants. **(A)** STAT5^-/-^ HeLa EpoR cells transduced with STAT5A-2Dto2A-meYFP or STAT5A-10Aand2D-meYFP were stimulated with 2 U/ml Epo as indicated. meYFP fluorescence was detected via confocal live cell imaging. w/o, without; scale bars: 50 µm. **(B)** STAT5^-/-^ HeLa EpoR cells transduced with STAT5A-WT-meYFP, STAT5A-2Dto2A-meYFP or STAT5A-10Aand2D-meYFP were stimulated with 1 U/ml Epo as indicated. Whole cell lysates (WCL) were analyzed by native PAGE (NP) or immunoblotting with the indicated antibodies. Note that the STAT5A-2Dto2A-meYFP sample was loaded twice. **(C)** STAT5^-/-^ HeLa EpoR cells transduced with STAT5A-WT-meYFP, STAT5A-Δ750-meYFP, STAT5A-2Dto2A-meYFP, STAT5A-10Aand2D-meYFP or STAT5A-Δ762-meYFP were stimulated with 2 U/ml Epo. Confocal images of meYFP fluorescence were acquired at 5-minute intervals. Images were segmented into nuclei and cytoplasm and fluorescence intensities within these compartments calculated using Arivis image analysis software. The diagram shows the ratio of fluorescence intensities in the nucleus and cytoplasm as a function of time. The y-axis is on a log_2_ scale.

To assess the effects of the deletion and point mutations on nucleocytoplasmic shuttling of STAT5A in a more quantitative manner, live-cell imaging experiments were performed in STAT5^-/-^ HeLa EpoR cells expressing selected STAT5A-meYFP mutants. Using imaging analysis software, cytoplasm and nuclei of cells were segmented, and the ratios of meYFP fluorescence calculated (Fig. 7C). The STAT5A mutants Δ750, 2Dto2A and 10Aand2D exhibited cytoplasmic retention with comparable kinetics, indicating that the point mutations cause complete inactivation of nuclear accumulation similar to the deletion mutant. In contrast, Δ762, the shortest deletion mutant capable of nuclear accumulation, was fully active with respect to nuclear accumulation. Interestingly, STAT5A-Δ762 translocated into the nucleus even faster than WT, which may be attributed to its smaller molecular size.

Next, we asked the question whether the acidic sequence motif must be present twice within the pSTAT5 dimer or if a single motif is sufficient to enable nuclear translocation. To address this question, we employed the recently published CATCHFIRE approach^32^. This method allows heterodimerization of two constructs when one protomer is tagged with FIREmate and the other with FIREtag. Heterodimerization is chemically induced by adding the fluorogenic substrate “match”, which fluoresces upon simultaneous binding to FIREmate and FIREtag. The corresponding STAT5A constructs were generated by fusing either FIREmate or FIREtag to the C-termini of both WT STAT5A (scheme in Fig. 8A) and STAT5A-2Dto2A, resulting in four distinct fusion proteins.

**Figure 8:**
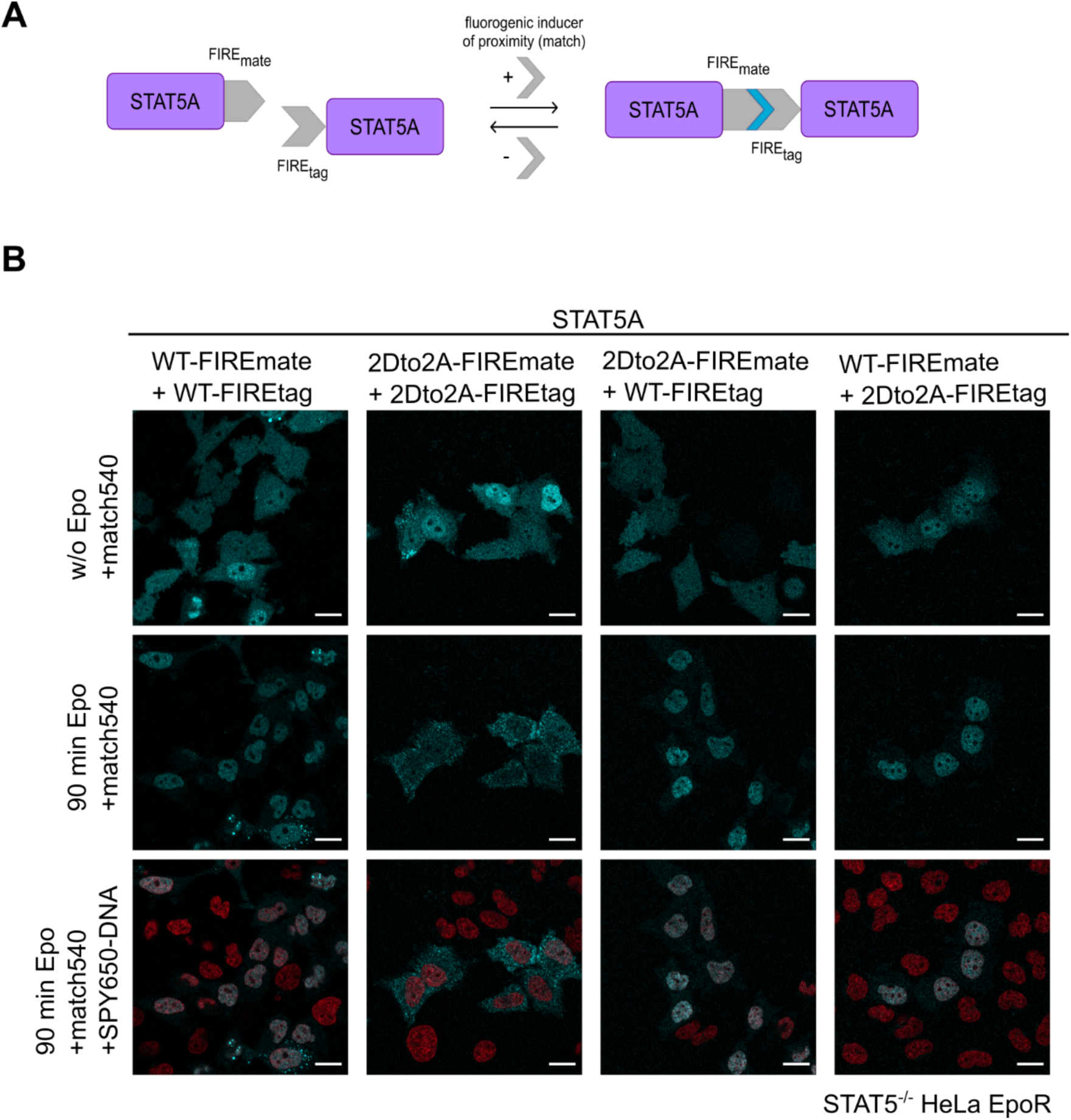
Subcellular localization of STAT5A-2Dto2A and STAT5A WT upon CATCHFIRE-mediated homo-and heterodimerization. **(A)** Schematic representation of the CATCHFIRE approach applied to STAT5A. Upon addition of match, the FIREmate and FIREtag constructs form dimers and match becomes fluorescent, indicated by the color change from grey to blue. **(B)** STAT5^-/-^ HeLa EpoR cells were transduced with a combination of FIREmate and FIREtag constructs of STAT5A-WT and STAT5A-2Dto2A as indicated. Nuclei were stained with SPY650-DNA. Cells were treated with 5 µM match540 for 15 min and subsequently with 2 U/ml Epo as indicated. SPY650-DNA (red) and match540 (cyan) fluorescence were detected via confocal live cell imaging. w/o, without; scale bars: 20 µm.

STAT5^-/-^ HeLa EpoR cells were transduced with all possible combinations of STAT5A WT and mutant FIREmate and FIREtag constructs. Dimerization was induced by the addition of match540 and monitored by live-cell imaging. Notably, match540-induced dimerization of STAT5A-FIREtag and STAT5A-FIREmate resulted in nuclear translocation of WT STAT5A homodimers only in a subset of cells. Consistent and complete nuclear import of STAT5A homodimers across all cells was achieved only upon addition of Epo (Fig. 8B, left images). The STAT5A-2Dto2A-FIREtag and STAT5A-2Dto2A-FIREtag constructs showed partial nuclear accumulation upon addition of match540, similar to STAT5A WT. Upon Epo stimulation, the match540-induced STAT5A-2Dto2A dimers exit the nucleus and accumulate in the cytoplasm, verifying our previous results obtained with the meYFP constructs (Fig. 8B, images second from the left). Remarkably, the heterodimerization of WT STAT5A with STAT5A-2Dto2A in both combinations of FIREmate and FIREtag resulted in nuclear accumulation of the heterodimers (Fig. 8B, images right and second from right). This finding clearly indicates that one functional TAD in a pSTAT5 dimer is sufficient for nuclear translocation.

## Discussion

STAT proteins are among the best studied transcription factors, but the precise molecular mechanisms governing their nucleocytoplasmic transport remain incompletely understood. In this study, we report an unexpected functional role of the transactivation domain (TAD) in the nuclear import of phosphorylated STAT5 (pSTAT5). STAT5 deletion mutants lacking the TAD do not enter the nucleus but accumulate in the cytoplasm while an analogous STAT3 deletion mutant is even more efficiently imported than WT (Fig. 1). The latter finding aligns with reports that the natural STAT3-β splice variant, which lacks the TAD, enters the nucleus more rapidly and is retained longer than full-length STAT3^33^. The aberrant cytoplasmic accumulation of STAT5-ΔTAD was observed across HeLa cells, HeLa cells lacking endogenous STAT5 and HEK cells, regardless of the meYFP-tag position or even in the absence of any tag (Fig. 2). The lack of nuclear import for N-terminally tagged STAT5A upon stimulation is consistent with the critical role of the N-terminal domain (NTD) in importing pSTAT dimers^17, 20, 21^. This is likely due to steric hindrance imposed by the tag on an essential function of the NTD in nuclear import. Accordingly, a STAT5A-ΔNTD mutant is not imported into the nucleus upon Epo stimulation (Suppl. Fig.6). Cytoplasmic accumulation of STAT5A-ΔTAD depends on phosphorylation at Y694 and dimerization (Fig. 3). Our results underscore a major role for the TAD in nuclear translocation of pSTAT5 dimers and support the concept that nuclear import requirements vary within the STAT family^5^, except for the essential function of the NTD. Notably, the steady-state distribution of STAT5A remains largely unchanged upon deletion of the TAD. Thus, the import of latent STAT5A is regulated by mechanisms distinct from those governing the nuclear import of phosphorylated STAT5A dimers. This observation has also been made for STAT1^9^, STAT3^21^, and the highly homologous STAT5B protein^17^.

As its name suggests, the TAD of STAT5 is indispensable for induction of target gene expression and has been extensively studied by others. The TAD is involved in various post-translational modifications, including acetylation, serine phosphorylation, SUMOylation and ubiquitination, yet no functional role in nuclear import has been established^34^. In gene induction, the interaction of the TAD with the histone acetylases (HAT) p300/CBP is most relevant^28^. Given that HATs acetylate not only histones but also other substrates such as STAT5^29, 35^, we investigated whether the reported acetylation sites K696 and K700 of STAT5A might control nuclear translocation. These lysines will most probably not be acetylated in the STAT5-ΔTAD mutant due to the loss of HAT interaction. Acetylation of STAT5A in lymphoid cells has been proposed to function as part of an acetylation/SUMOylation switch. Acetylation of K696 is essential for dimerization as well as transcriptional activity and is blocked by SUMOylation of the same residue^29^. Interestingly, blue-native PAGE analysis demonstrated that Epo-induced STAT5 dimerization is increased rather than abolished by mutation of these critical lysines. Correspondingly, the STAT5A-K696R/K700R-meYFP mutant accumulates in the nucleus upon Epo stimulation (Suppl. Fig. 3). Thus, acetylation of K696 or K700 is not essential for STAT5 activation or nuclear import.

A stretch of 12 amino acids (F751-H762) within the TAD has previously been studied in relation to transactivation^30^ and turnover of pSTAT5^31^. Chen et al. identified this region as critical for the ubiquitination of nuclear pSTAT5 by mediating its interaction with an unidentified E3 ligase^27^. Therefore, deletion of these amino acids is expected to stabilize STAT5, resulting in accumulation of nuclear pSTAT5 rather than its degradation. Consistent with this, treatment with the proteasome inhibitor MG132 did not prevent the disappearance of STAT5A-ΔTAD from the nucleus upon Epo stimulation (Suppl. Fig. 2B). Furthermore, our immunoblot analyses showed that TAD deletion had no major impact on the levels of pSTAT5 or total STAT5. Hence, we conclude that TAD deletion primarily affects nucleocytoplasmic shuttling rather than protein stability. The same 12 amino acids are also essential for the transactivation function of STAT5, with particular importance attributed to the hydrophobic residues F751 and L753^30^. Consequently, we examined the role of the F751-H762 region and found it to be indispensable for nuclear import of pSTAT5 (Fig. 5). Remarkably, fusion of these 12 amino acids to STAT5A-ΔTAD was sufficient to restore nuclear import (Fig. 6). However, not the hydrophobic residues but the acidic residues D754 and D758 are essential for nuclear import (Fig. 7A and Suppl. Fig. 4). Quantitative analysis of imaging data revealed that the nuclear import function of the TAD was entirely lost upon mutation of these two residues (Fig. 7C). Thus, within this small segment of the protein, the structural determinants for transactivation and nuclear import can be distinguished. Our data suggests that F751-H762 contributes to a dimer-specific NLS (dsNLS) unique to STAT5. This dsNLS is also required for nuclear translocation of STAT5 upon phosphorylation by the oncogenic kinase BCR-ABL (Fig. 2B). Given the essential role of STAT5 in chronic myeloid leukemia and other cancers^34^, targeting its dsNLS may represent a promising therapeutic strategy.

We utilized the recently developed CATCHFIRE approach to perform chemically induced heterodimerization of STAT5 variants with a fluorogenic inducer^32^. With this approach, we demonstrated that a single intact TAD is sufficient for the nuclear import of pSTAT5 dimers (Fig. 8). This finding raises the question of how, on one hand, dimerization of pSTAT5 is a prerequisite for nuclear translocation while, on the other hand, a single critical dsNLS motif is sufficient. We hypothesize that the dsNLS motif in the TAD is complemented by an additional dsNLS motif located elsewhere in the pSTAT5 dimer, together forming a functional dsNLS.

The C-terminal region of STAT5A, including the TAD, undergoes serine phosphorylation, with phosphorylation at S725 and S779 playing critical roles in leukemogenic transformation^36^ and with S779 being required for nuclear translocation during oncogene-induced activation^37^. Notably, S779 is absent in the STAT5A-Δ762 mutant, which still retains competence for Epo-induced nuclear import (Fig. 5). Thus, phosphorylation of S779 is not generally required for nuclear import of pSTAT5A. Another serine residue, S756, is located within the 12 amino acids stretch essential for nuclear import. Mutation of S756 to alanine did not affect nuclear import of pSTAT5A (Suppl. Fig. 5). Thus, activity of the STAT5-specific dsNLS is not regulated by serine phosphorylation.

Interestingly, the TAD has been implicated in Crm1-mediated nuclear export of STAT1^38^. Upon inhibition of Crm1-dependent nuclear export by leptomycin B, STAT5A-ΔTAD accumulates slightly in the nucleus, similar to WT STAT5. This indicates that the TAD is not essential for steady-state nuclear export of STAT5. Nuclear depletion upon stimulation still occurs even in the presence of leptomycin B, demonstrating that the cytoplasmic accumulation of pSTAT5A-ΔTAD is not due to enhanced Crm1-mediated nuclear export. Instead, we interpret that pSTAT5A-ΔTAD is retained in the cytoplasm because it is defective in nuclear import (Fig. 2D). Steady-state nuclear export of non-phosphorylated STAT5A-ΔTAD is still ongoing and once exported STAT5A-ΔTAD becomes phosphorylated in the cytoplasm, it adds to the trapped cytoplasmic pool of pSTAT5A-TAD. Over time, this leads to the observed nuclear clearance and cytoplasmic accumulation.

Taken together, our findings reveal an unexpected and critical role of the TAD in the nuclear import of pSTAT5 dimers. The TAD uniquely contributes to a dimer-specific nuclear localization signal (dsNLS) that, together with complementary motifs within the dimer, facilitates efficient nuclear translocation. This mechanism distinguishes nuclear import of phosphorylated dimers from nucleocytoplasmic shuttling of latent monomers. Furthermore, the dependence of STAT5 nuclear import on this dsNLS, which is also engaged during oncogenic activation by BCR-ABL, underscores its potential as a precise therapeutic target in myeloid leukemias and related cancers. Future studies dissecting this import machinery may offer novel strategies to modulate STAT5 activity in disease.

## Methods

### Antibodies

Anti-pY694-STAT5 (#9351, 1:1000, WB, Cell Signaling, USA), anti-pY694-STAT5-XP (#4322, 1:200, IF, Cell Signaling, USA), anti-pY705-STAT3 (#9131, 1:1000, WB, Cell Signaling, USA), anti-STAT5A (A7, sc-166479, 1:1000; Santa Cruz Biotechnology, USA) and anti-GFP (600-101-215, 1:1000, WB, Rockland, USA) were used as indicated. Secondary horseradish peroxidase (HRP)-conjugated anti-rabbit-IgG (P044801-2, 1:2000), anti-mouse-IgG (P044701-2, 1:2000) and anti-goat-IgG (P016002-2, 1:2000) were purchased from Dako (Denmark). Secondary fluorescent antibodies anti-rabbit-IgG Alexa Fluor 488 (A11034, 1:1000, IF), anti-rabbit-IgG Alexa Fluor 647 (A31573, 1:2000, WB), anti-mouse-IgG Alexa Fluor 647 (A31571, 1:2000, WB) and anti-goat-IgG Alexa Fluor 488 (A11055, 1:2000, WB) were purchased from Thermo Fisher Scientific (USA).

### Cell culture

HeLa T-REx^®^ (Invitrogen, USA) and HEK293T cells were cultured in Dulbeccós Modified Eagle Medium GlutaMAX I (DMEM, Gibco, USA) supplemented with 10% fetal calf serum (FCS, Capricorn Scientific, Germany) and 1% penicillin/streptomycin (10.000 U/ml penicillin, 10 mg/ml streptomycin, Sigma-Aldrich, USA), in a humidified atmosphere at 37°C with 5% CO_2_. Stably transfected HeLa T-REx^®^ (Invitrogen) cell lines were selected with 200 µg/ml hygromycin (InvivoGen, France) and 15 µg/ml blasticidin (Invivogen). HeLa T-REx^®^ cell lines were treated with 5 ng/ml doxycycline (Sigma-Aldrich) for 24 h, to induce the expression of HA-EpoR, using the Flp-In system (Invitrogen).

### Lentiviral mediated transduction of cells

For constitutive expression of STAT(-meYFP) constructs in (STAT5^-/-^) HeLa EpoR cells, a lentiviral gene delivery system was used to transduce the cells with the cDNA encoding for the target gene. All work with lentivirus was performed under guidelines of the RNAi consortium (S2 safety conditions). The cDNA of interest was cloned into the lentiviral pLentiLox expression vector (Vector Core, University of Michigan, USA). On day one, HEK293T packaging cells were seeded and the next day, HEK293T cells were co-transfected with psPAX2 (packaging plasmid, Vector Core), pCMV-VSV-G (envelope plasmid, Vector Core) and the lentiviral vector containing the GOI (gene of interest). For high transfection efficiency, the TransIT^®^-LT1 (Mirus, USA) transfection reagent was used, according to manufacturer’s instructions. The day after transfection, the medium of the HEK293T cells was replaced by high serum growth medium containing 30% FCS. The virus-containing supernatant from the transfected HEK293T cells was collected 24 h after medium change, supplemented with 8 µg/ml polyprene (Sigma-Aldrich), added to previously seeded (STAT5^-/-^) Hela EpoR cells and incubated overnight.

### Establishing STAT5-/- HeLa EpoR knockout cells using CRISPR/Cas9

HeLa T-REx HA-EpoR cells (generated as described in Fahrenkamp et al.^26^) were transfected with plasmids for CRISPR/Cas9 gene editing using the TransIT^®^-LT1 transfection reagent according to the manufacturer’s instructions. First, the targeted chromosomal fragment of STAT5A was deleted using four pX335A plasmids (MPI, Münster, Germany) encoding for paired sgRNAs targeting exon 4 and intron 4 of the STAT5A gene (ST5A-1sp/rp - ST5A-4sp/rp, sequences see below). A day after transfection, the medium was exchanged by selection medium containing 1 µg/ml puromycin and 250 µg/ml hygromycin. The next day, cells were washed once with PBS, detached from the plate with 0.5 ml Trypsin-EDTA and counted. To obtain single cell clones, cells were either seeded at a density of 0.5 cells per well onto 96-well plates or at a density of 50 cells per 10 cm plate. Dishes or plates were incubated and checked for colonies. The grown colonies were picked and transferred to 24-well plates. Genomic DNA was isolated from the colonies and parental cells using the QIAamp® DNA Mini Kit (QIAGEN, Germany) according to manufacturer’s instructions. A PCR with primers flanking the sequence to be deleted by Cas9 activity was performed (sequences of flanking primers see below). Positive clones were further analyzed for protein expression by Western blot. In case of successful knockout of STAT5A, STAT5A knockout (STAT5^+/-^) HeLa T-REx HA-EpoR cells were transiently transfected with four sgRNA/Cas9 expressing pX335A plasmids (MPI, Münster) encoding for paired sgRNAs targeting exon 2 and intron 2 of the STAT5B gene (ST5B-1sp/rp - ST5B-4sp/rp, sequences see below). The deletion and subsequent analysis of STAT5B knockout cells (sequences of flanking primers see below) were carried out using the same procedures as described for the generation and characterization of STAT5A knockout cells.

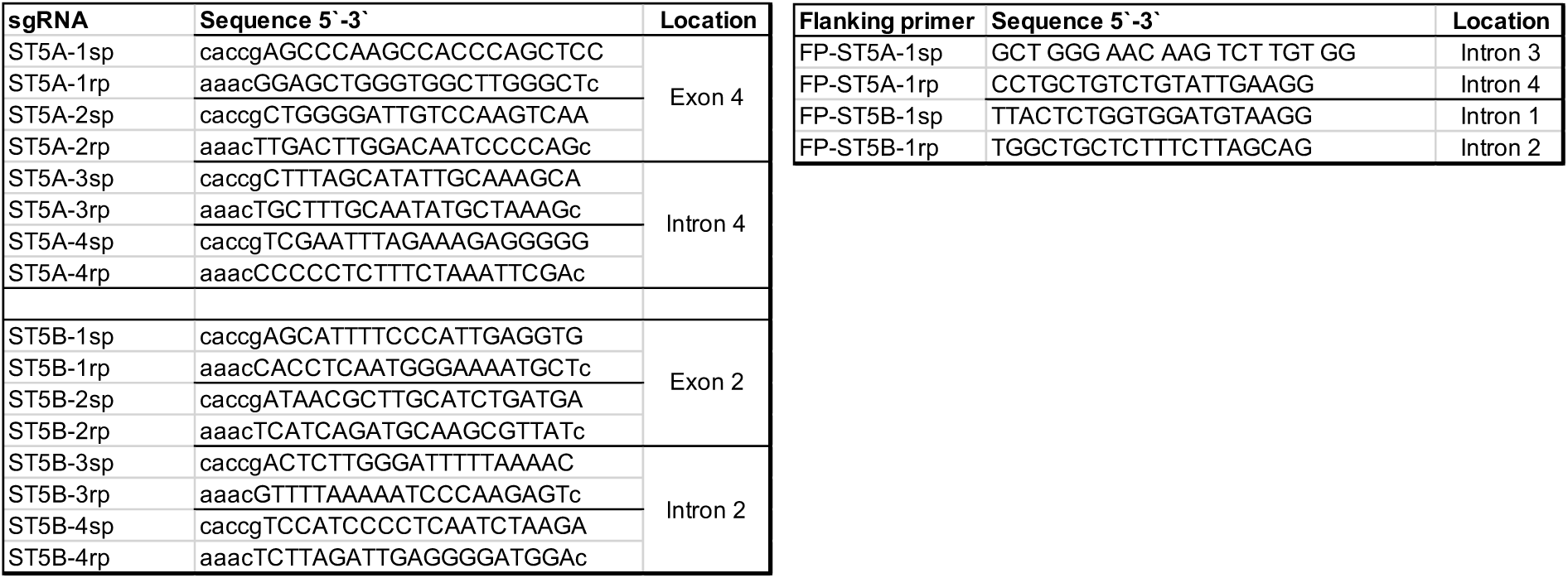

### Cell lysates, SDS-PAGE and immunoblotting

Cells were lysed using RIPA lysis buffer (50 mM Tris-HCl pH 7.4, 150 mM NaCl, 1 mM EDTA, 1 mM NaF, 0.5% (v/v) Nonidet P-40, 15% (v/v) Glycerol), supplemented with phosphatase inhibitor Na_3_VO_4_ (1 M) and protease inhibitors PMSF (0.25 mM), Aprotinin (5 µg/ml) and Leupeptin (5 µg/ml). Cell lysates were loaded onto a 10% SDS-gel, proteins separated and blotted onto a PVDF membrane by electrophoresis. Immunodetection was performed with specific primary antibodies followed by HRP-conjugated or fluorescently labeled secondary antibodies. Proteins were visualized using enhanced chemiluminescence (ECL) solution supplemented with 0.03% (w/v) H_2_O_2_ for 1 min at RT and detected with a LAS-4000 luminescence imaging device (Fujifilm, Japan). Proteins labeled with fluorescent secondary antibodies were detected with the Azure c600 imaging system (Azure Biosystems, USA).

### Native PAGE

The NativePAGE^TM^ Novex® Bis-Tris Gel System (Invitrogen) was used according to manufacturer’s instructions. Protein lysates were prepared and a total amount of 25 µg WCL, adjusted to an equal volume of 25 µl using NP sample buffer, supplemented with 2% Digitonin (Sigma-Aldrich), were loaded. Prior to loading, 0.5 µl NativePAGE^TM^ 5% G-250 Sample Additive (Invitrogen) were added to the samples, mixed and centrifuged at 16,000x g for 1 min at 4°C. The samples were loaded onto NativePAGE™ 4-16% Bis-Tris Protein Gels (1.0 mm) and run at 50 V over night or at 150 V for 2.5 h at 4°C. The electrophoresis was performed in a XCell SureLockTM MiniCell chamber (Invitrogen). The outer chamber was filled with 1x concentrated NP running buffer (Invitrogen) and the inner chamber with NP running buffer supplemented with 0.5% (v/v) of the 20x NativePAGE^TM^ Cathode Additive (Invitrogen). The wet gel, containing meYFP fluorescence labelled fusion proteins, was analyzed using the Azure c600 imaging system (Azure Biosystems).

### Glyoxal fixation and immunostaining

Cells were seeded at a density of 1x10^5^ cells per well in 12-well plates directly on sterile 18 mm glass coverslips in 1 ml culture medium. Expression of the inducible GOI was induced by addition of 5 ng/ml doxycycline for 24 h. Prior to fixation, doxycycline treated cells were stimulated with 1 U/ml Epo (Immunotools, Germany) for 60 min. The culture medium was removed, cells were washed two times with 500 µl PBS and fixed with 500 µl glyoxal solution (pH 5.0) for 30 min on ice, followed by 30 min at RT. After washing two times with PBS, the cells were treated with 500 µl of quenching buffer (PBS with 1 mM MgCl_2_, 0.1 CaCl_2_ and 50 mM NH_4_Cl) for 20 min at RT, to attenuate autofluorescence. The cells were washed once with PBS, followed by permeabilization and blocking of the samples with 500 µl 0.1% Triton-X-100 in PBS supplemented with 2.5% BSA for 15 min at RT. Primary and secondary antibodies were each diluted in 0.1% Triton-X-100 in PBS supplemented with 1% BSA. First the primary antibody was incubated for 60 min at RT, followed by a washing step and incubation of the fluorescently labeled secondary antibody for 60 min at RT. Cells were washed three times with PBS, once with ddH_2_O and the immunostained coverslips were mounted on microscope slides using Immu-Mount (Thermo Scientific, USA) mounting medium. Until image acquisition by confocal microscopy, the slides were stored protected from light at 4°C.

### Live cell imaging

For live cell imaging analysis, 1.5x10^4^-2x20^4^ cells were seeded onto µ-Slide 8 Well IbiTreat chambers (Ibidi, Germany) in 250 µL cell culture medium 48 h prior to imaging. One day after seeding, the expression of the inducible GOI (e.g EpoR) was induced by addition of 5 ng/ml doxycycline for 24 h. Images were acquired with a Zeiss LSM710 or Zeiss LSM980 with Airyscan 2 confocal laser scanning microscope (Zeiss, Germany) equipped with an incubation chamber (PeCon, Germany). Cells were imaged at 37°C and 5% CO_2_, using a LD C-Apochromat 40x/1.1 W Korr M27 or LD LCI Plan-Apochromat 40x/1.2 Imm Korr DIC M27 objective (Zeiss).

#### CATCHFIRE

FIREmate and FIREtag plasmids were purchased from The Twinkle Factory (France). STAT5^-/-^ HeLa EpoR cells were lentivirally transduced with STAT5A-FIREmate and STAT5A-FIREtag constructs (WT and/or the 2Dto2A-mutant). The experiment was designed as described in paragraph Live cell imaging. Nuclei were stained with SPY650-DNA (Spirochrome, Switzerland (SC501); 1:2000) for 30 min. Imaging was carried out using the Airyscan 2 detector of an inverted Zeiss LSM 980 confocal laser scanning microscope . Image acquisition was performed using an LD LCI Plan-Apochromat 40x/1.2 Imm Korr DIC M27 objective (water immersion) and CO-8Y mode. Images were taken with a pixel size of 0.099 µm and a pixel dwell time of 0.32 µs. SPY650-DNA was excited using a 639 nm diode laser, while match540 (ChemScene; USA) was excited with a 488 nm diode laser. To induce dimerization of STAT5A, double-transduced cells co-expressing STAT5A-FIREmate and STAT5A-FIREtag were required. 5 µM match540 were added and double-transduced cells identified by the appearance of fluorescence. After 15 min of incubation with match540, images of the unstimulated cells were acquired. Subsequently, cells were stimulated with 2 U/ml Epo, and images were taken every 15 min over a 90 min period. All images were processed using the *Airyscan Processing* module in automated mode within the Zeiss ZEN Blue software 3.12.

### Live cell imaging for quantification of nucleus/cytoplasm ratio

The experiment was conducted as outlined in the Live cell imaging section. Multiple positions within each well each containing different STAT5A mutant cell lines, were identified and imaged. Prior imaging, nuclei were stained with SPY650-DNA (Spirochrome (SC501); 1:2000) for 30 min. Following cell stimulation with 2 U/ml Epo, time-lapse imaging was performed at 5-minute intervals for a total duration of 120 minutes, employing the Definite Focus 3 system for focus stabilization and automated image acquisition. Images were taken with a commercial inverted Zeiss LSM 980 confocal laser scanning microscope equipped with an Airyscan 2 detector. Image acquisition was performed using a Plan-Apochromat 10x/0.45 M27 objective (Zeiss), employing the LSM Plus mode. Images were taken with a pixel size of 0.18 µm and a pixel dwell time of 0.34 µs. SPY650-DNA was excited using a 639 nm diode laser, while meYFP was excited with a 488 nm diode laser. All images were processed using the *LSM Plus* module in automated mode within the Zeiss ZEN Blue software 3.8.

### Quantification of nucleus/cytoplasm ratio with Arivis

Fluorescence intensity measurements were performed on segmented image datasets to quantify spatial distribution within the cells. Fluorescence images were analyzed using Arivis Pro software (Zeiss, version 4.1.1). The mean fluorescence intensity was extracted separately from the nuclear and cytoplasmic regions, and a nuclear-to-cytoplasmic (N/C) fluorescence intensity ratio was computed to describe subcellular signal partitioning. All downstream statistical analyses of N/C intensity ratios, as well as generation of graphs, were carried out using Excel (Microsoft, USA).

### Intensity profile

To analyze co-localization and LMB mediated nuclear accumulation of STAT5-meYFP, intensity values were extracted from images using the profile function of the ZEN 2012 software (Zeiss). A diagram showing the intensity profile was created using the GraphPad Prism 8 software (GraphPad Software, USA).

## Acknowledgements

This study was funded by grants to G.M.N. from the German Research Foundation (Deutsche Forschungsgemeinschaft, DFG; project number 417911533) as part of the Clinical Research Unit CRU 344. This work was supported by the Confocal Microscopy Facility, a Core Facility of the Interdisciplinary Center for Clinical Research (IZKF) Aachen within the Faculty of Medicine at RWTH Aachen University. The authors would like to thank Hildegard Schmitz-van de Leur for plasmid cloning.

## Author contributions

Experiments: S.E., M.B.M., A.K. and H.S.; Writing original draft: S.E. and G.M.N.; Review and editing: M.B.M., A.K. and H.S.; Conceptualization and funding acquisition: G.M.N.

