## Supplemental Figures for "Unexpected functional role of the transactivation domain for nuclear import of STAT5"

**Supplementary Figures**

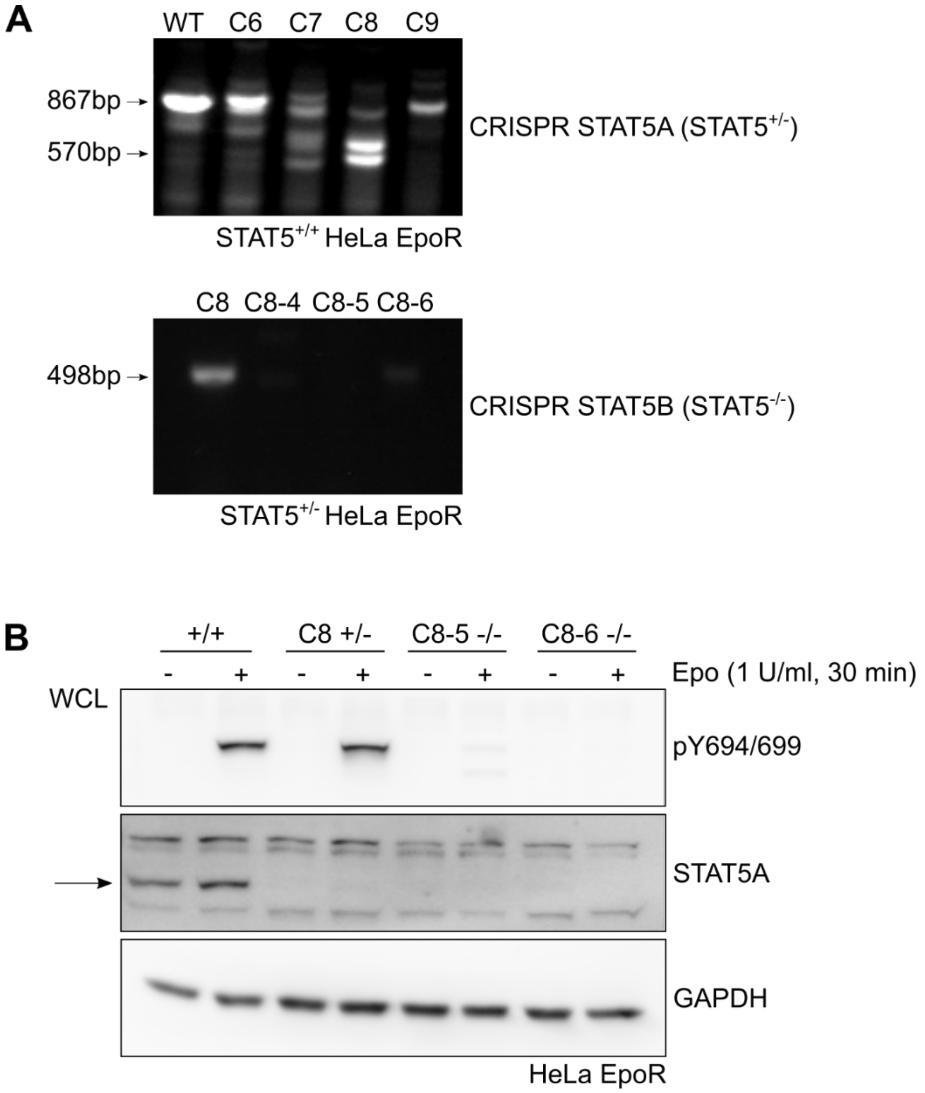


**Suppl. Fig. 1: Generation of STAT5$^{-/-}$ HeLa EpoR cells by CRISPR/Cas9-mediated gene targeting of STAT5A and STAT5B.**

**(A)** Upper panel: A defined deletion in the STAT5A gene was introduced into STAT5$^{+/+}$ HeLa T-REx HA-EpoR cells by transfection with four sgRNA/Cas9 expression plasmids (pIST5A 1-4). The presence of the intended genomic deletions was assessed by PCR using a STAT5A-specific primer pair (FP-ST5A-fwd1/rev1), followed by size analysis of the amplicons by agarose gel electrophoresis. The wild-type STAT5A locus yielded an 867 bp amplicon, whereas the deletion allele produced a shorter fragment of 570 bp, corresponding to the expected loss of 297 bp.

Lower panel: A targeted deletion in the STAT5B gene was introduced in STAT5$^{+/-}$ HeLa T-REx HA-EpoR clone C8 cells using four sgRNA/Cas9 expression plasmids (pIST5B 1–4). Detection of deletions was performed by PCR amplification with the specific primer pair FP-ST5B-fwd1/rev1, followed by agarose gel electrophoresis analysis. The expected wild-type STAT5B amplicon was 498 bp in size, while mutant amplicons representing deletions were not detected under these conditions.

**(B**) Expression of STAT5A and Epo-induced phosphorylation of STAT5A and STAT5B were analyzed in HeLa T-REx HA-EpoR cells (STAT5$^{+/+}$), STAT5A-deficient clone 8 (C8+/-) and two STAT5A/B-deficient subclones (C8-5 -/- and C8-6 -/-). Cells were stimulated with Epo as indicated. Whole cell lysates (WCL) were prepared and subjected to Western blot analysis. Phosphorylated STAT5A and STAT5B were detected using antibodies against phospho-STAT5 (pY694/699), while total STAT5A levels were assessed with a STAT5A-specific antibody. GAPDH served as a loading control.

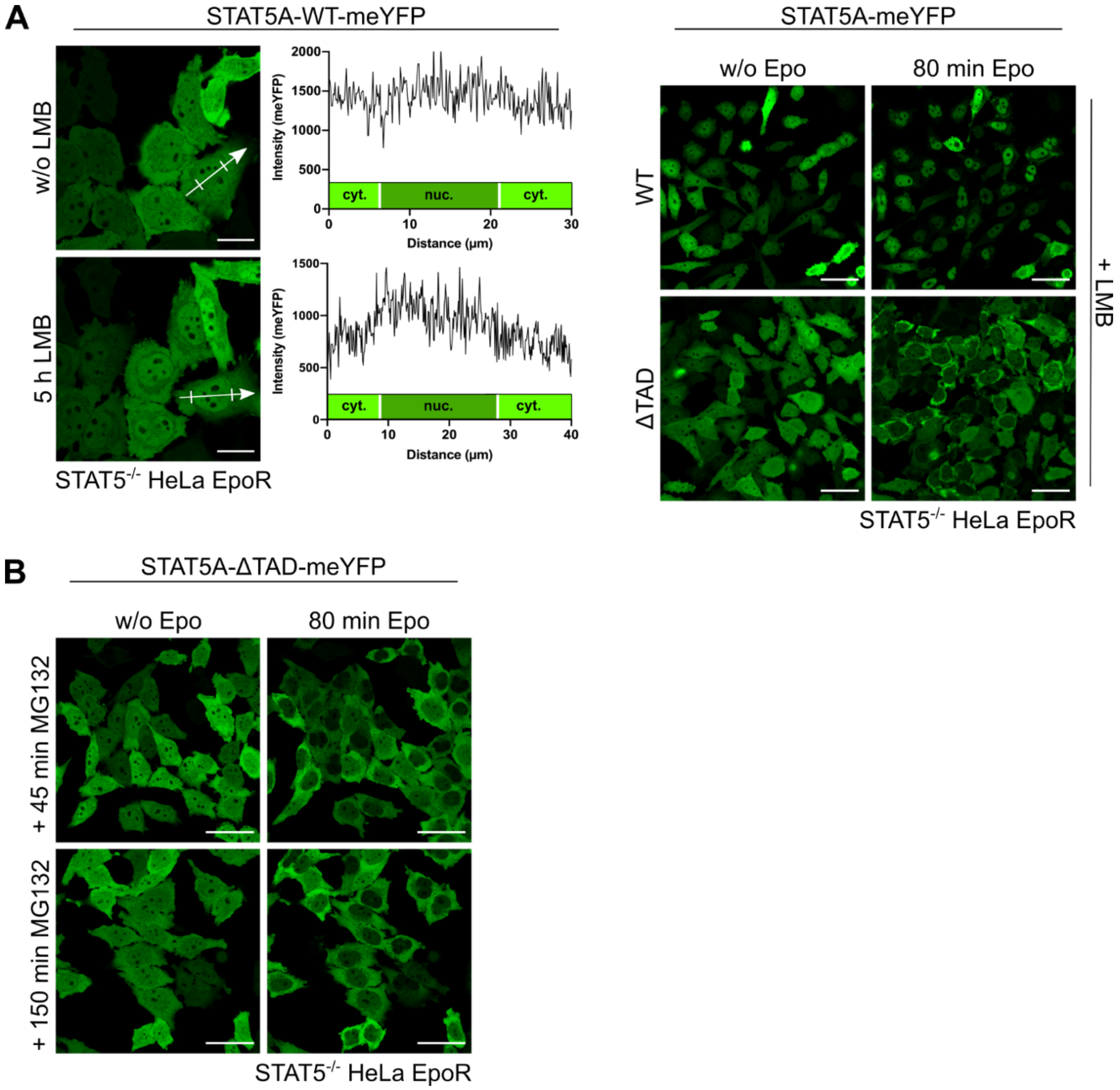


**Suppl. Fig. 2: Nucleocytoplasmic shuttling of STAT5A WT and ΔTAD deletion mutant after inhibition of CRM1-mediated nuclear export or MG132-mediated proteasome inhibition.**

**(A)** $STAT5^{-/-}$ HeLa EpoR cells transduced with STAT5A-WT-meYFP were treated with 30 nM leptomycin B (LMB) as indicated. Left panel: meYFP fluorescence was detected via confocal live cell imaging. Scale bars: 20 µm. Intensity profiles of meYFP fluorescence were generated along the white arrows depicted in the images, with cytoplasm (cyt.)/nucleus (nuc.) boundaries indicated. Right panel: $STAT5^{-/-}$ HeLa EpoR cells transduced with STAT5A-WT-meYFP or STAT5A-ΔTAD-meYFP were preincubated with 30 nM LMB and subsequently stimulated with 1 U/ml Epo as indicated. meYFP fluorescence was detected via confocal live cell imaging. Scale bars: 50 µm.

**(B)** $STAT5^{-/-}$ HeLa EpoR cells transduced with STAT5A-ΔTAD-meYFP were preincubated with 40 µM MG132 and subsequently stimulated with 1 U/ml Epo as indicated. meYFP fluorescence was detected via confocal live cell imaging. w/o, without Epo; scale bars: 50 µm.

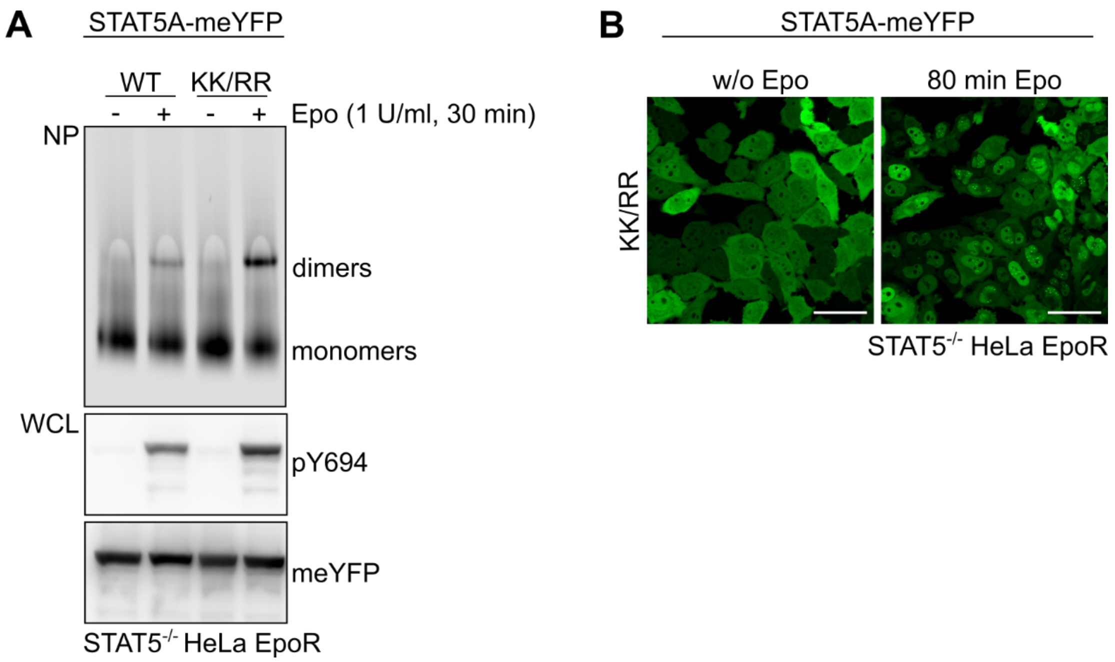


**Suppl. Fig. 3: Dimerization and subcellular localization of the STAT5A K696R/K700R mutant.**

**(A)** STAT5$^{-/-}$ HeLa EpoR cells transduced with STAT5A-WT-meYFP or STAT5A-K696R/K700R-meYFP (KK/RR) were stimulated with Epo as indicated. Whole cell lysates (WCL) were analyzed by native PAGE (NP) or immunoblotting with the indicated antibodies.

**(B)** STAT5$^{-/-}$ HeLa EpoR cells transduced with STAT5A-K696R/K700R-meYFP (KK/RR) were stimulated with 1 U/ml Epo as indicated. meYFP fluorescence was detected via confocal live cell imaging. w/o, without; scale bars: 50 µm.

| STAT5A | Amino acid sequence | | | Localization before/after Epo stimulation | |
|---|---|---|---|---|---|
| | 1-750 | …751-762… | …793… | - | + |
| WT<br>-(meYFP) | M1-E750 | FDLDESMDVARH | 793-STOP<br>793-meYFP-STOP | 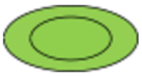 | 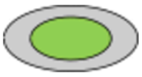 |
| ΔTAD<br>-(meYFP) | M1-A717 | ---- | STOP<br>meYFP-STOP | 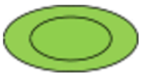 | 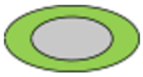 |
| Δ750<br>-meYFP | M1-E750 | ---- | meYFP-STOP | 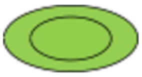 | 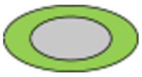 |
| Δ756<br>-meYFP | M1-E756 | FDLDES | meYFP-STOP | 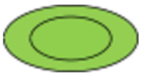 | 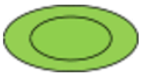 |
| Δ762<br>-meYFP | M1-E750 | FDLDESMDVARH | meYFP-STOP | 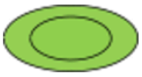 | 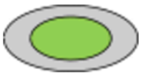 |
| ΔTAD+12aa<br>-meYFP | M1-A717 | FDLDESMDVARH | meYFP-STOP | 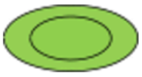 | 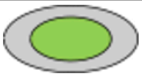 |
| 12A<br>-meYFP | M1-E750 | AAAAAAAAAAAA | 793-meYFP-STOP | 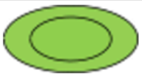 | 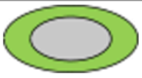 |
| 9WT3A<br>-meYFP | M1-E750 | FDLDESMDVAAA | 793-meYFP-STOP | 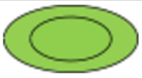 | 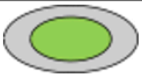 |
| 3A9WT<br>-meYFP | M1-E750 | AAADESMDVARH | 793-meYFP-STOP | 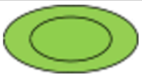 | 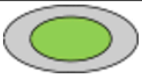 |
| 3A6WT3A<br>-meYFP | M1-E750 | AAADESMDVAAA | 793-meYFP-STOP | 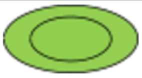 | 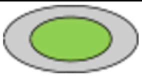 |
| 3WT6A3WT<br>-meYFP | M1-E750 | FDLAAAAAAARH | 793-meYFP-STOP | 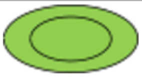 | 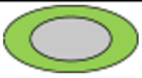 |
| 2Dto2A<br>-meYFP | M1-E750 | FDLAESMAVARH | 793-meYFP-STOP | 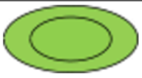 | 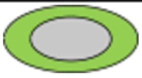 |
| 10Aand2D<br>-meYFP | M1-E750 | AAADAAADAAAA | 793-meYFP-STOP | 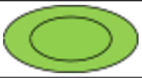 | 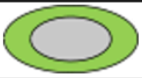 |

**Suppl. Fig. 4: Overview of STAT5A deletion and alanine-scanning constructs and schematic representation of their subcellular localizations.**

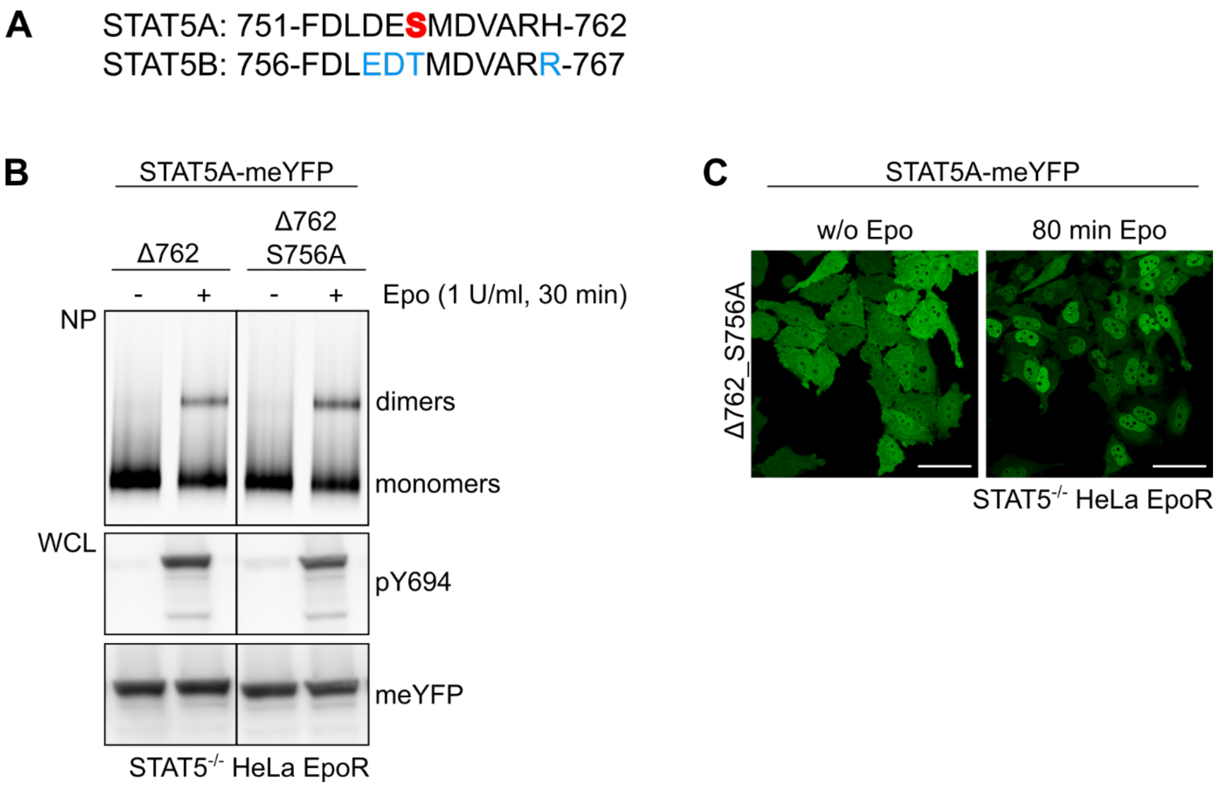


**Suppl. Fig. 5: Dimerization and subcellular localization of the STAT5A Δ762_S756A mutant.**

**(A)** Amino acid sequences of STAT5A (residues Phe751 to His762) and STAT5B (residues Phe756 to Arg767) are shown. Serine 756 in STAT5A is highlighted in red. Conserved amino acid substitutions in STAT5B compared to STAT5A are indicated in blue.

**(B)** STAT5$^{-/-}$ HeLa EpoR cells transduced with STAT5A-Δ762-meYFP or STAT5A-Δ762_S756A-meYFP were stimulated with Epo as indicated. Whole cell lysates (WCL) were analyzed by native PAGE (NP) or immunoblotting with the indicated antibodies.

**(C)** STAT5$^{-/-}$ HeLa EpoR cells transduced with STAT5A-Δ762_S756A-meYFP were stimulated with 1 U/ml Epo as indicated. meYFP fluorescence was detected via confocal live cell imaging. w/o, without; scale bars: 50 µm.

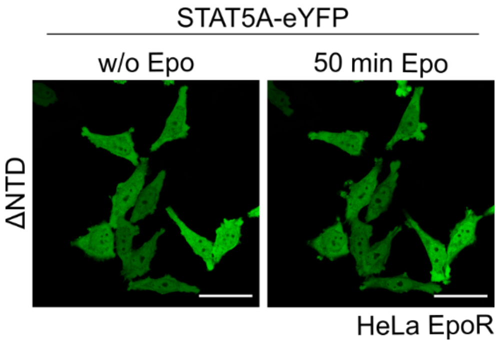


**Suppl. Fig. 6: Subcellular localization of the STAT5A ΔNTD deletion mutant.**
HeLa EpoR cells transduced with STAT5A-ΔNTD-eYFP were stimulated with 1 U/ml Epo as indicated. eYFP fluorescence was detected via confocal live cell imaging. w/o, without; scale bars: 50 µm.
